# A Bayesian model for human directional localization of broadband static sound sources

**DOI:** 10.1101/2022.10.25.513770

**Authors:** Roberto Barumerli, Piotr Majdak, Michele Geronazzo, Federico Avanzini, David Meijer, Robert Baumgartner

**Affiliations:** Acoustics Research Institute, Austrian Academy of Sciences, 1040 Vienna, Austria; Dept. of Management and Engineering, University of Padova, Italy; Dyson School of Design Engineering, Imperial College London, London, United Kingdom; Dept. of Computer Science, University of Milano, Italy

## Abstract

Humans estimate sound-source directions by combining prior beliefs with sensory evidence. Prior beliefs represent statistical knowledge about the environment while sensory evidence is acquired from auditory features such as interaural disparities and monaural spectral shapes. Models of directional sound localization often impose constraints on the contribution of these features to either the horizontal or vertical dimension. Instead, we propose a Bayesian model that more flexibly incorporates each feature according to its spatial precision and integrates prior beliefs in the inference process. We applied the model to directional localization of a single, broadband, stationary sound source presented to a static human listener in an anechoic environment. We simplified interaural features to be broadband and compared two model variants, each considering a different type of monaural spectral features: magnitude profiles and gradient profiles. Both model variants were fitted to the baseline performance of five listeners and evaluated on the effects of localizing with non-individual head-related transfer functions (HRTFs) and sounds with rippled spectrum. The model variant with spectral gradient profiles outperformed other localization models. This model variant appears particularly useful for the evaluation of HRTFs and may serve as a basis for future extensions towards modeling dynamic listening conditions.

## I. INTRODUCTION

When localizing a sound source, human listeners have to deal with numerous sources of uncertainty [1]. Uncertainties originate from ambiguities in the acoustic signal encoding the source position [2] as well as the limited precision of the auditory system in decoding the received acoustic information [3, 4]. Bayesian inference describes a statistically optimal solution to deal with such uncertainties in the process of perceptual decision making [5] and has been applied to model sound localization in various ways [6–9].

Common approaches of sound localization models rely on the evaluation of several spatial auditory features. Head-related transfer functions (HRTFs) describe all the spatially dependent acoustic filtering produced by the listener’s ears, head, and body [10] and have been used to derive spatial auditory features. The way to quantify or extract those features is a matter of debate. In particular, a large variety of monaural spectral-shape features have been studied [11–17], with spectral magnitude profiles [14, 17] and spectral gradient profiles [12, 15] being the most established ones. Despite such details, there is consensus that the interaural time and level differences (ITDs and ILDs) [1] as well as some form of monaural spectral shapes are important features for the directional localization of broadband sound sources [18].

In order to decode the spatial direction from the auditory features, models rely on the assumption that listeners have learned to associate acoustic features with spatial directions [13, 19]. In fact, the interaural features are particularly informative about the horizontal dimension [1] though rather ambiguous with respect to the vertical dimension, where evidence from the monaural spectral features is more important [12]. This anisotropic relevance of different features is the reason for why specific auditory features are often studied along a single dimension of the so-called modified interaural-polar coordinate system [20], with the lateral angle along the horizontal left/right dimension and the polar angle along the vertical and front/back dimension. However, this geometric separation is a simplification. Monaural spectral features, for instance, can also contribute to the inference process in the direction estimation along the lateral dimension [21–23]. Hence, directional sound localization models should rather exploit the joint information encoded by all auditory features.

Such joint information has already been considered in a model of directional sound localization based on Bayesian inference [6]. This model computes a spatial likelihood function from a precision-weighted integration of a set of noisy acoustic features. Then, the perceived source direction is assumed to be at the maximum of that likelihood function. While this model was built to assess which spatial information can be accessible to the auditory system, its predictions overestimate the actual human performance yielding unrealistically low front-back confusion rates and localization errors [24]. Still, in order to model human performance, this model can serve as a solid basis for improvements such as the consideration of monaural spectral features, the integration of response noise involved in typical localization tasks, and the incorporation of prior beliefs.

Prior beliefs are important in the process of Bayesian inference because they reflect the listener’s statistical knowledge about the environment, helping to compensate for uncertainties in the sensory evidence [25]. For example, listeners seem to effectively increase precision in a frontal localization task by assuming source directions to be more likely located at the eye-level rather than at extreme vertical positions [8]. However, such an increase in precision may come at the cost of decreasing accuracy. As it seems, the optimal accuracy-precision trade-off in directional localization depends on the statistical distribution of sound sources [26]. While listeners seem to adjust their prior beliefs to changes in the sound-source distribution [26, 27], they may also establish long-term priors reflecting the distribution of sound sources in their everyday environment.

Here, we introduce a Bayesian inference model to predict the performance of a listener in estimating the direction of static broadband sounds. Similar to [6], our model implements a noisy feature extraction and probabilistically combines interaural and monaural spatial features. These features are then compared with templates of spatial features, obtained from listener-specific HRTFs, to generate the sensory evidence in the form of a likelihood function. Subsequently, the sensory evidence is combined with prior beliefs emphasizing directions at the eye level [8]. The estimated source direction is selected from the resulting (posterior) spatial representation according to a Bayesian decision function. In a final stage, the model incorporates response scattering [15] to account for the uncertainty introduced by pointing responses in localization experiments.

For evaluation, we considered a model variant based on spectral amplitudes and a model variant based on spectral gradients [15]. Each model’s free parameters were fitted to the sound-localization performance of individual listeners [28]. We then tested the simulated responses of both model variants against human responses from sound-localization experiments investigating the effects of non-individual HRTFs [29] and ripples in the source spectrum [30].

The paper is organized as follows: Sec. II describes the auditory model (Sec. II A) and explains the parameter estimation (Sec. II B). Then, Sec.III evaluates the model’s performance by comparing its estimations to the actual performance of human listeners. Finally, Sec. IV discusses the model’s relevance as well as its limitations, and outlines its potential for future extensions.

## II. METHODS

### A. Model description

The proposed auditory model consists of three main stages, as shown in Fig. 1: 1) The feature extraction stage determines the encoded acoustic spatial information represented as a set of spatial features altered by noise; 2) The Bayesian inference integrates the sensory evidence resulting from the decoding procedure based on feature templates with the prior belief and forms a perceptual decision, and 3) The response stage transforms the perceptual decision in a directional response by corrupting the estimation with uncertainty in the pointing action.

**FIG. 1:**
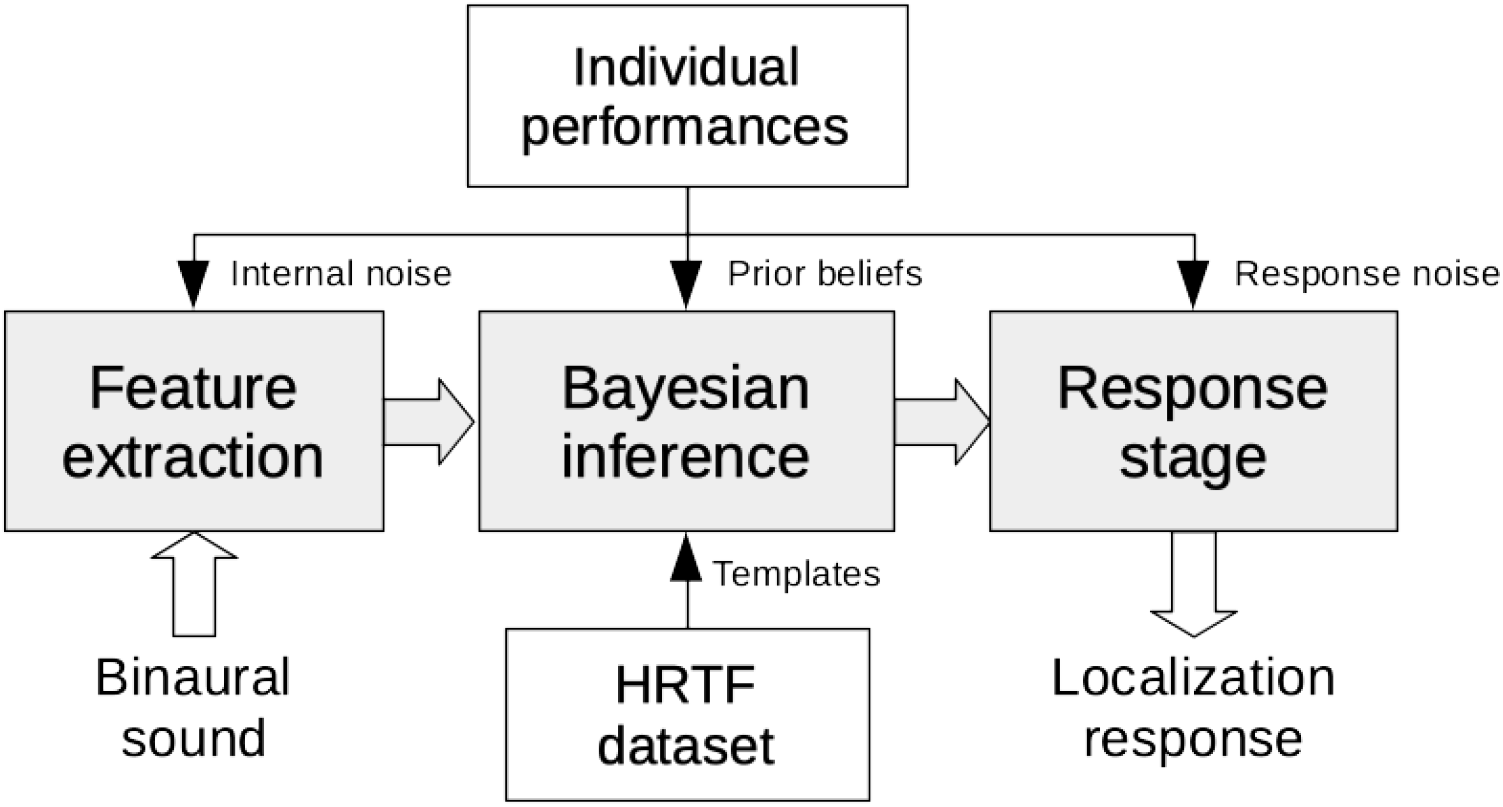
Model structure. Gray blocks: Model’s processing pipeline consisting of 1) the *feature extraction* stage to compute spatial features from the binaural sound; 2) the *Bayesian inference* stage integrating the sensory evidence obtained by comparison with feature templates and with prior beliefs in order to estimate the most probable direction; and 3) the *response stage* transforming the internal estimate to the final localization response. White blocks: Elements required to fit the model to an individual subject consisting of listener performances in estimating sound direction and individual HRTF dataset.

#### 1. Feature extraction

The spatial auditory features are extracted from the sensory input which is provided by the directional transfer function transformed in the time domain (i.e. the HRTF processed to remove the direction independent component [31]). We follow [6] in that we decode the spatial information provided by a single sound source via the binaural stimulus from a vector of spatial features:

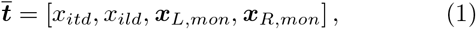

where *x*_*itd*_ denotes a scalar ITD feature, *x*_*ild*_ a scalar ILD feature, and a vector that concatenates monaural spectral features for left ear, ***x***_*L,mon*_, and right ear, ***x***_*R,mon*._ Each feature is assumed to be extracted by different neural pathways responsible to deliver encoded spatial information to higher levels of the auditory system [1, 4].

Assuming broadband and spatially stationary sources, interaural features can be heavily approximated by means of wideband estimators [18, 32]. The ILD was approximated as the time-averaged broadband level difference between the left and right channels [18]. The ITD was estimated by first processing each channel of the binaural signal with a low-pass Butterworth filter (10^*th*^ order and cutoff 3000 Hz) and an envelope extraction step based on the Hilbert transform. Then, the ITD value is computed with the interaural cross-correlation method which is a good estimator of perceived lateralization in static scenarios with noise bursts [32]. In addition, we applied the transformation proposed by Reijniers *et al*. [6] to compensate the increasing uncertainty levels for increasing ITDs [33] resulting in a dimensionless quantity with a more isotropic variance:

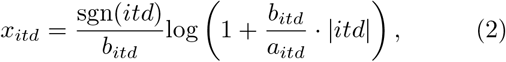

with *itd* denoting ITDs in *μs* and the parameters *a*_*itd*_ = 32.5*μs* and *b*_*itd*_ = 0.095 and ‘sgn’ indicating the sign function (for details on the derivation based on signal detection theory, see Supplementary Information from [6]). An example of the interaural features as functions of the lateral angle is shown in Fig. 2.

**FIG. 2:**
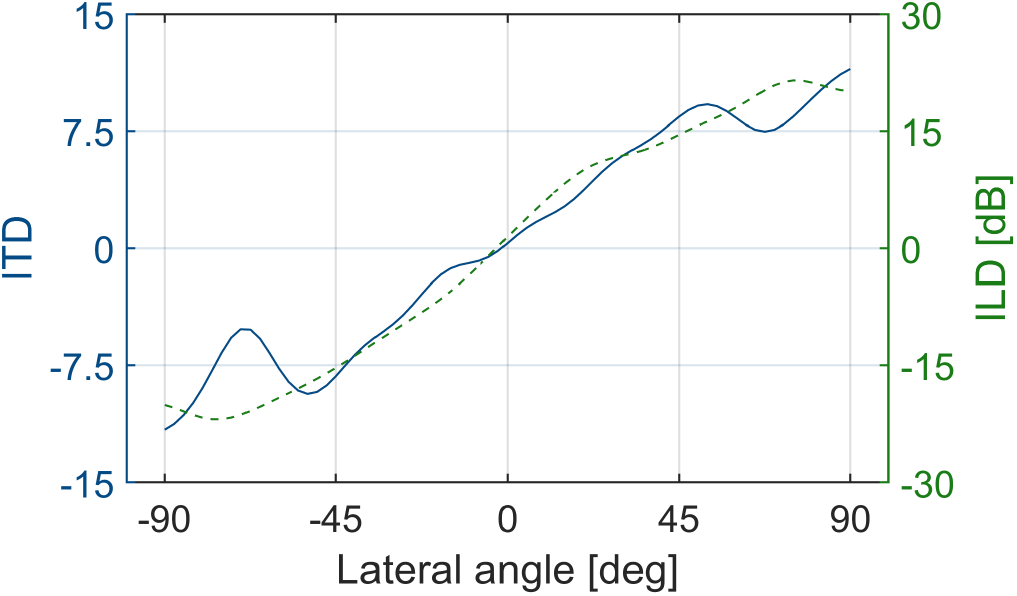
Interaural features as functions of lateral angle in the horizontal frontal plane (*ϕ* = 0°). Left axis (blue solid line): Transformed ITD *x*_*itd*_ (dimensionless), see Eq. 2. Right axis (green dashed line): ILD (in dB) obtained from the magnitude profiles. Example for subject NH12 [28].

Monaural spectral features, ***x***_*{L,R}, mon*_, were derived from approximate neural excitation patterns. To approximate the spectral resolution of the human cochlea, we processed the binaural signal by the gammatone filter-bank with non-overlapping equivalent rectangular bandwidths [34, 35], resulting in *N*_*B*_ = 27 bands within the interval [0.7, 18] kHz [36, 37]. Followed by half-wave rectification and square-root compression to model hair-cell transduction [e.g., 38, 39], it results in the unit-less excitation:

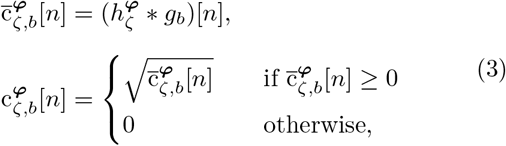

where subscripts *ζ* ∈ {*L, R*} indicate the left and right ears, *n* = 1, …, *N* is the time index, *b* = 1, …, *N*_*B*_ is the band index, *g*_*b*_[*n*] is the corresponding gammatone filter and 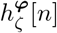 is the binaural signal in a normalized scale with sound direction ***φ*** (i.e. a pair of head-related impulse responses or their convolution with a source signal).

We thus define the spectral feature for the magnitude profiles (MPs) with the vector ***x***_*ζ,MP*_. This vector is the collection of root mean square amplitudes across time in decibels for each of the spectral bands for each ear:

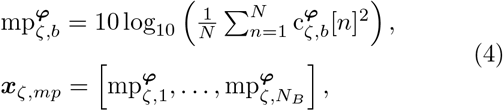

where the function 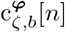is defined in Eq. 3.

An alternative spectral feature can be computed by positive gradient extraction over the frequency dimension. It has previously been shown that integrating such spectral features in an auditory model provides good agreement with human localization performance [15]. Therefore, we define a second possible spectral feature based on gradient profiles (GPs) with the vector ***x***_*ζ,GP*_. It includes the gradient extraction as an additional processing step:

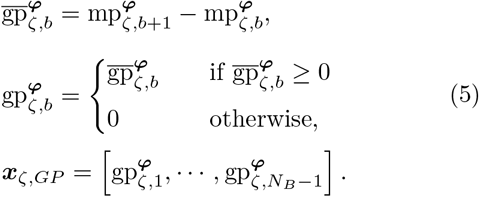

A visualization of these monaural features is shown in Fig. 3.

**FIG. 3:**
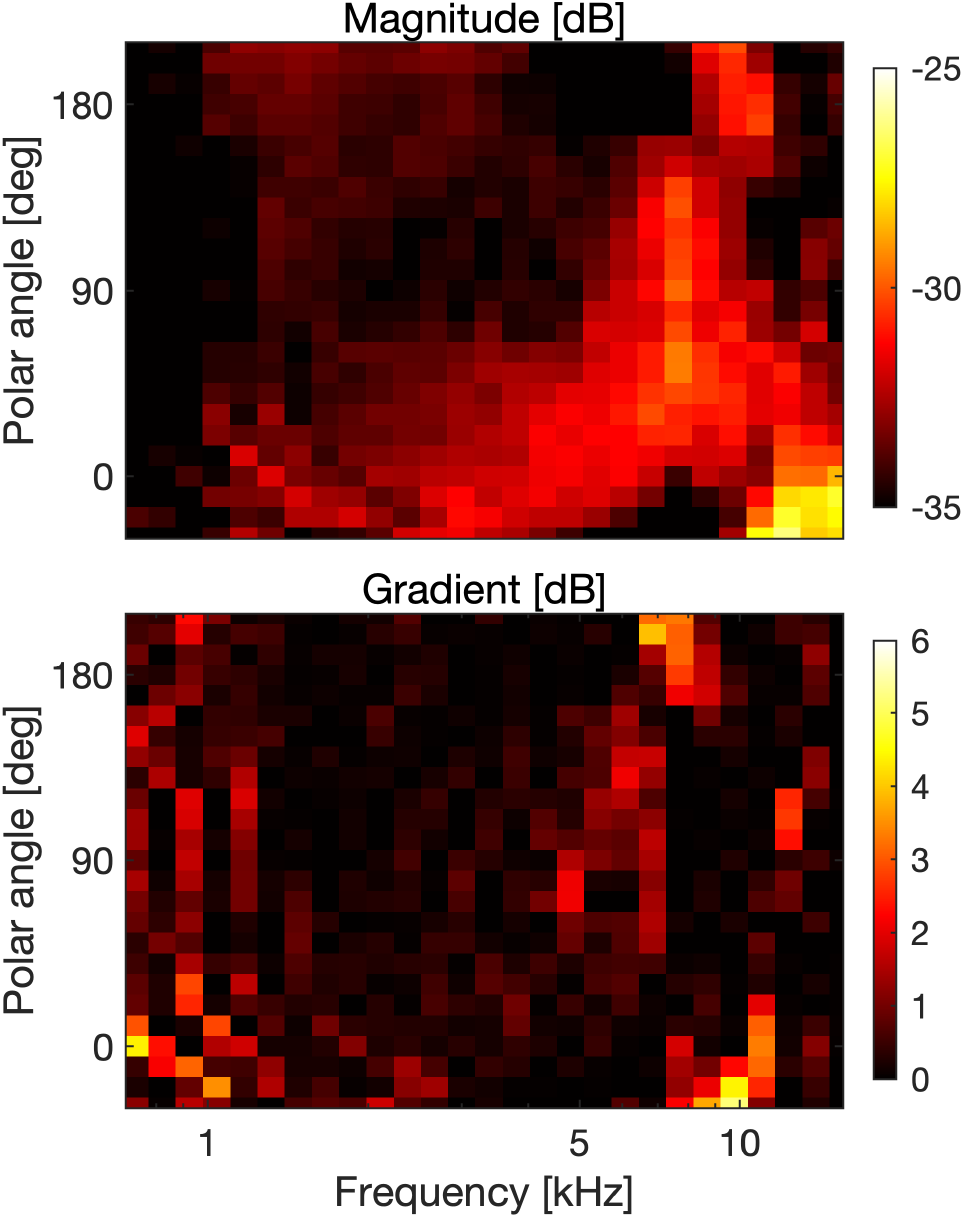
Monaural spectral features as a function of polar angle in the median plane (*θ* = 0°). Top: Features obtained from the magnitude profiles. Bottom: Features obtained from the gradient profiles. Example for the left ear of subject NH12 [28].

To demonstrate the impact of monaural spectral feature type, we will analyze the results of both variants with the corresponding feature spaces defined as follows:

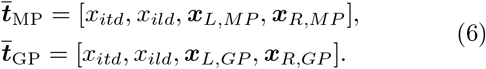

Limited precision in the feature extraction process leads to corruption of the features and can be modelled as additive internal noise [6]. Hence, we define the noisy internal representation of the target features as:

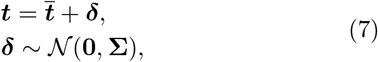

where **Σ** is the covariance matrix of the multivariate Gaussian noise. Furthermore, we assume each spatial feature to be processed independently and thus to be also corrupted by independent noise [1]. Hence, the covariance matrix **Σ** is defined as:

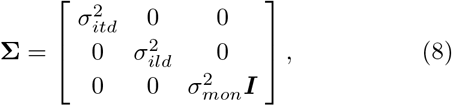

with 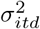 and 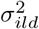 being the variances associated with the ITDs and ILDs and 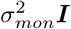 being the covariance matrix for the monaural features where ***I*** is the identity matrix and the scalar *σ*_*mon*_ represents a constant and identical uncertainty for all frequency bands.

#### 2. Bayesian inference

The observer infers the sound direction ***φ*** from the spatial features in ***t*** while taking into account potential prior beliefs about the sound direction. Within the Bayesian inference framework [5], this requires to weight the likelihood *p*(***t*** | ***φ***) with the prior *p*(***φ***) to obtain the posterior distribution by means of Bayes’ law as:

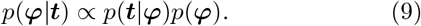

The likelihood function is implemented by comparing ***t*** with the feature templates. Similarly to [6], the template ***T*** (***φ***) contains noiseless features of Eq. 1 for every sound direction ***φ***. While the sound direction is defined on a continuous support, our implementation sampled it over a uniform spherical grid with a spacing of 4.5°between points (*N*_*φ*_ = 1500 over the full sphere). Template features were computed from the listener-specific HRTFs. To accommodate non-uniform HRTF acquisition grids, we performed spatial interpolation based on spherical harmonics with order *N*_*SH*_ = 15, followed by Tikhonov regularization [40].

Since the templates are constructed without noise there exists a one-to-one mapping between direction and template features. This allows us to write the likelihood function for each point of the direction grid as:

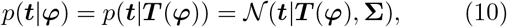

where **Σ** represents the learned precision of the auditory system (i.e. the sensory uncertainty ***δ*** reported in Eq. 7). Finally, we interpret the a-priori probability *p*(***φ***) to reflect long-term expectations of listeners where prior probabilities are modelled as uniformly distributed along the horizontal dimension but centered towards the horizon as [8]. In particular, we extend the results from Ege *et al*. for sources positioned in the front and as well as back positions with:

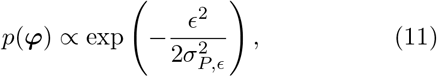

with *ϵ* denoting the elevation angle of ***φ*** and 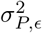 the variance of the prior distribution [8]. For simplicity, the prior definition is based for the spherical coordinate system. Importantly, the origin of that prior is currently unknown and its implications are discussed in Sec. IV.

According to Eq. 9, a posterior spatial probability distribution is computed for every sound by optimally combining sensory evidence with prior knowledge [25]. As shown in Fig. 4, the most probable direction of the source ***φ*** is then selected as the maximum a-posteriori (MAP) estimate:

**FIG. 4:**
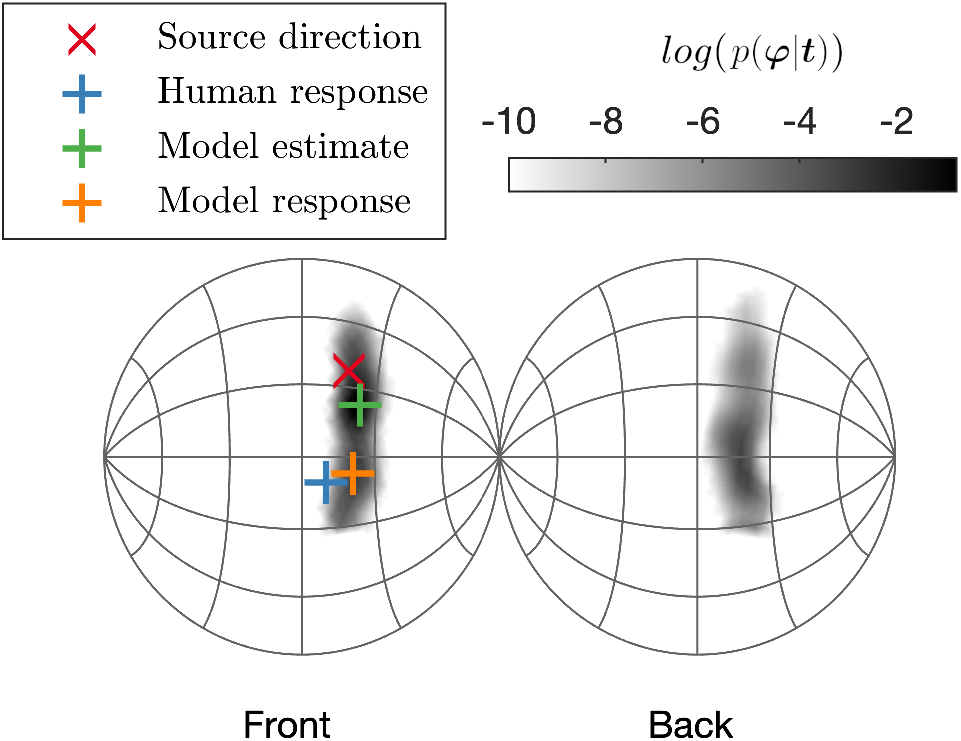
Example of the model estimating the direction of a broadband sound source. Red: Actual direction of the sound source. The grayscale represents the posterior probability distribution *p*(φ|t), shown, in order to increase the readability, on a logarithmic scale. Green: Direction inferred by the Bayesian-inference stage (without the response stage). Orange: Direction inferred by the model (with the response stage). Blue: Actual response of the subject.

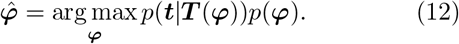

#### 3. Response stage

After a sound direction estimate has been inferred, experiments usually require the listener to provide a motor response (e.g. manual pointing). To account for the uncertainty introduced by such responses, we incorporate post-decision noise in the model’s response stage. Following the approach from previous work [15], we distort the location estimate by additive, direction-independent (i.e. isotropic noise) Gaussian noise:

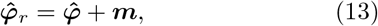

where ***m*** ∼ vMF(0, *κ*_*m*_) is a von-Mises-Fisher distribution with zero mean and concentration parameter *κ*_*m*_. The concentration parameter *κ*_*m*_ can be interpreted as a standard deviation 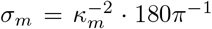 [deg]. The contribution of the response noise is also visible in Fig. 4, where the final estimate is scattered independently of the spatial information provided by the a-posteriori distribution. With Eq. 13, the model outputs the response of the estimated sound source direction.

### B. Parameter estimation

The model includes the following free parameters: *σ*_*ild*_, *σ*_*mon*_ (amount of noise per feature; *σ*_*itd*_ was fixed to 0.569 as in [6]), *σ*_*P,ϵ*_ (directional prior), and *σ*_*m*_ (amount of response noise). Because of the structure of the model, these parameters jointly contribute to the prediction of performance in both lateral and polar dimensions. To roughly account for listener-specific differences in localization performance [2], the parameters were fitted to match individual listener performance.

As for the objective fitting function, we selected a set of performance metrics widely used in the analysis of behavioral localization experiments [28, 29, 41], for a summary see [42]. A commonly used set of metrics contains the quadrant error rate (QE, i.e., frequency of polar errors larger than 90°), local polar errors (PE, i.e., root mean square error in the polar dimension that are smaller than 90° limited to lateral angles in the range of ±30°), and lateral errors (LE, i.e., root mean square error in the lateral dimension) [29]. We accounted for the inherent stochasticity of the model estimations by averaging the simulated performance metrics over 300 repetitions of the *N*_*φ*_ = 1550 directions in the HRTF dataset (i.e. Monte Carlo approximation with 465000 model simulations). Model parameters were jointly adjusted in an iterative procedure (see below) until the relative residual between the actual performance metric *E*_*a*_ and the predicted performance metric *E*_*p*_ was minimized below a metric-specific threshold *τ*_*E*_, i.e.,

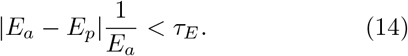

We set the thresholds to *τ*_*LE*_ = 0.1, *τ*_*P E*_ = 0.15, and *τ*_*QE*_ = 0.2 because those values were feasible for all subjects. In addition, the QE was transformed with the rationalized arcsine function to handle small and large values adequately [43].

We ran the estimation procedure separately for each feature space in Eq. 6 and each listener. First, initial values of the parameters were derived from previous literature: the variance of the prior distribution was set to *σ*_*P,ϵ*_ = 11.5°as in [8]. The interaural feature noise was set to *σ*_*ild*_ = 1 dB, reflecting the range of ILD thresholds for pure tones [44]. The starting value for the monaural feature noise was set to *σ*_*mon*_ = 3.5 dB similarly to in [6]. The response noise standard deviation was set to *σ*_*m*_ = 17°as the sensorimotor scatter found in [15]. Second, in an iterative procedure, *σ*_*m*_ was optimized to minimize the residual error relative to the PE metric and, similarly, *σ*_*mon*_ was adjusted to match the QE metric. Then, *σ*_*ild*_ was decreased to reach the LE metric. These steps were reiterated until the residual errors between actual and simulated metrics was less than the respective threshold. This procedure limited the *σ*_*m*_ to the interval [5°, 20 °] and used a step-size of 0.1° *σ*_*mon*_ was defined in the interval [0.5, 10] dB with a step-size of 0.05 dB; *σ*_*ild*_ was defined in the interval of [0.5, 2] dB with a step-size of 0.5 dB. If the procedure did not converge, we decreased *σ*_*P,ϵ*_ by 0.5°and reattempted the parameter optimization procedure.

## III. RESULTS

We first report the quality of model fits to the calibration data itself [28] in Sec. III A. Then, Sec. III B quantitatively evaluates the simulated performances of our two model variants and of two previously proposed models against data from two additional sound localization experiments.

### A. Parameter fits

The parameter estimation procedure was done with both model variants, based on either 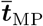 or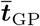, and for five individuals tested in a previous study [28]. In that experiment, naive listeners were asked to localize broadband noise bursts of 500 ms duration presented from various directions on the sphere via binaural rendering through headphones based on listener-specific directional transfer functions. The subjects were wearing a headmounted display and were asked to orient the pointer in their right hand to the perceived sound-source direction. The fitting procedure converged for both models and all subjects. Notably, subject NH15 required to reduce the step size of *σ*_*m*_ to 0.1°to meet the convergence criteria. Tab. I reports the estimated parameters *σ*_*m*_, *σ*_*P,ϵ*_, *σ*_*mon*_ and *σ*_*ild*_ for every listener. The amount of response noise was similar for both model types. Tab. II contrasts the predicted performance metrics with the actual ones, averaged across listeners.

**TABLE I:** Parameters estimated to fit the models to actual subjects’ performance [28] for both model variants where either magnitude profiles (MPs) or gradient profiles (GPs) were the monaural spectral features.

| Variant | Subject | $\sigma_{P,\epsilon}$ [deg] | $\sigma_{ild}$ [dB] | $\sigma_{mon}$ [deg] | $\sigma_m$ [deg] |
| --- | --- | --- | --- | --- | --- |
| MP | NH12 | 11.50 | 0.50 | 3.40 | 8.50 |
|  | NH15 | 10.00 | 0.50 | 3.20 | 14.27 |
|  | NH16 | 11.50 | 1.00 | 3.60 | 11.00 |
|  | NH17 | 11.50 | 0.50 | 4.10 | 14.30 |
|  | NH18 | 11.50 | 1.00 | 6.50 | 14.00 |
| GP | NH12 | 11.50 | 0.50 | 1.10 | 8.50 |
|  | NH15 | 11.00 | 0.50 | 1.25 | 14.30 |
|  | NH16 | 11.50 | 1.00 | 1.25 | 11.50 |
|  | NH17 | 11.50 | 1.00 | 1.60 | 14.00 |
|  | NH18 | 11.50 | 1.00 | 2.10 | 15.00 |

**TABLE II:** Predicted performance metrics averaged across all subjects and directions (±1 standard deviation across subjects) for both model variants. Actual data from [28].

| Metric | Actual | Predicted |  |
| --- | --- | --- | --- |
|  |  | MP | GP |
| LE [deg] | $12.25 \pm 2.43$ | $12.97 \pm 2.50$ | $13.18 \pm 2.66$ |
| PE [deg] | $32.73 \pm 3.44$ | $31.20 \pm 4.04$ | $29.78 \pm 4.01$ |
| QE [%] | $7.83 \pm 7.11$ | $8.32 \pm 5.75$ | $9.80 \pm 5.23$ |

More in detail, Fig. 5 compares predicted localization performance to the actual performance of subjects estimating the direction of a noise burst for different spherical segments [28]. The predicted LEs and PEs, both as functions of the actual lateral and polar angles, respectively, were in good agreement with those from the actual experiment. Instead, the simulated QE metric failed to mimic the front back asymmetries present in four subjects. Finally, only small differences were observed between the two feature spaces 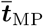and 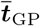.

**FIG. 5:**
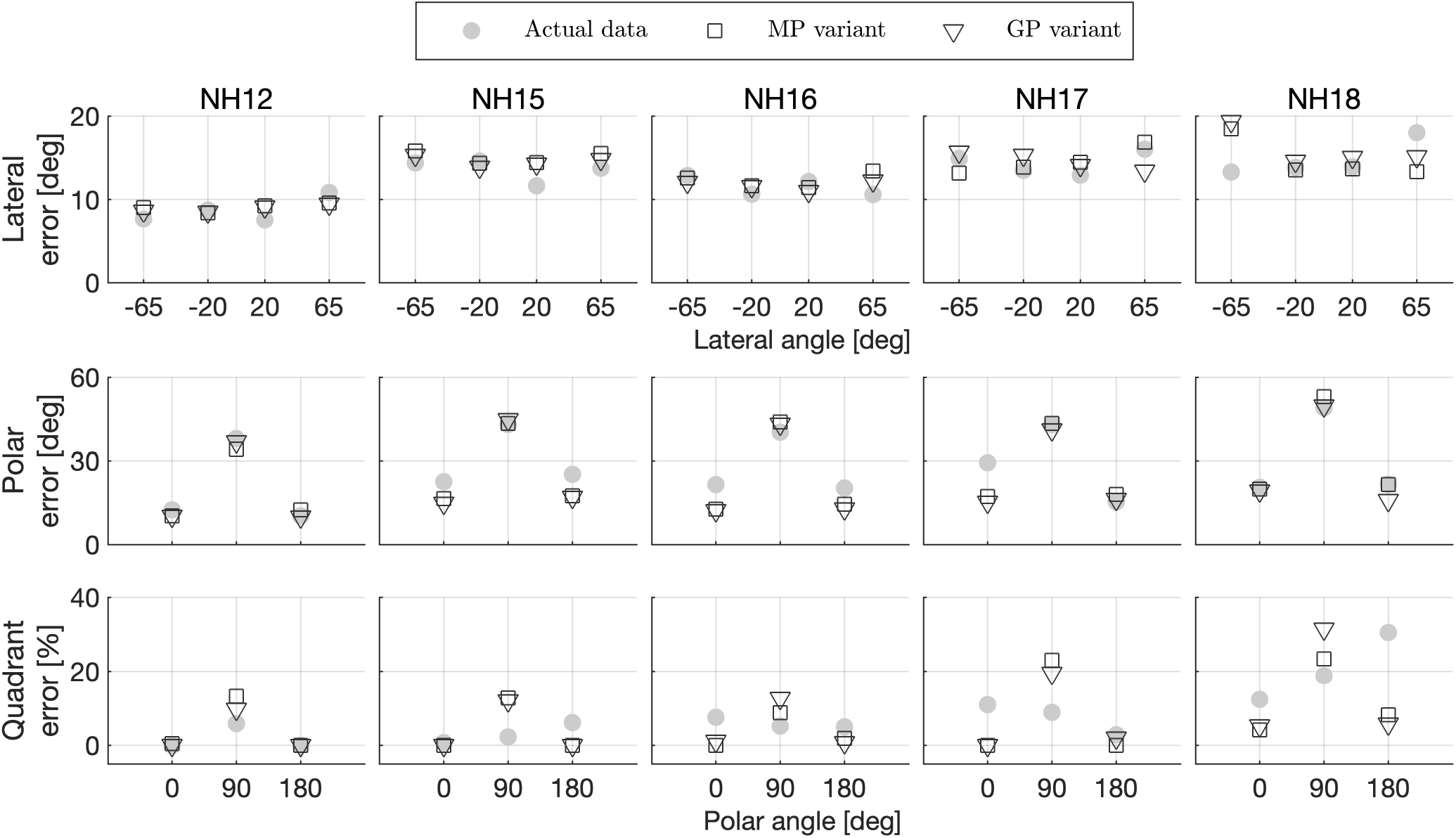
Sound-localization performance as functions of sound-source direction. Open symbols: Predictions obtained by the two model’s variants based on either spectral magnitude profiles (MPs) or gradient profiles (GPs). Filled grey symbol: Actual data from [28]. Top row: Lateral error, calculated for all targets with lateral angles of −65°*±* 25° −20°*±* 20° 20°*±* 20° and 65°*±* 25°. Center and bottom rows: Polar error and quadrant error rates, respectively, calculated for all median-plane (*±*30°) targets with polar angles of 0°*±* 30° 90°*±* 60° and 180°*±* 40°.

#### Contribution of model stages

Fig. 6 illustrates the effects of different model stages on target-specific predictions. The example shows direction estimations from subject NH16 localizing broadband noise bursts [28] and the corresponding predictions of the model based on 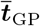 with different configurations of priors and response noise: without both (a), with priors only (b), and with both (c). While adding response noise scatters the estimated directions equally across spatial dimensions (compare c to b), including the spatial prior only affects the polar dimension (compare b to a). As observed in the actual responses, the prior causes more of the simulated estimations to be biased towards the horizon (0°and 180°).

**FIG. 6:**
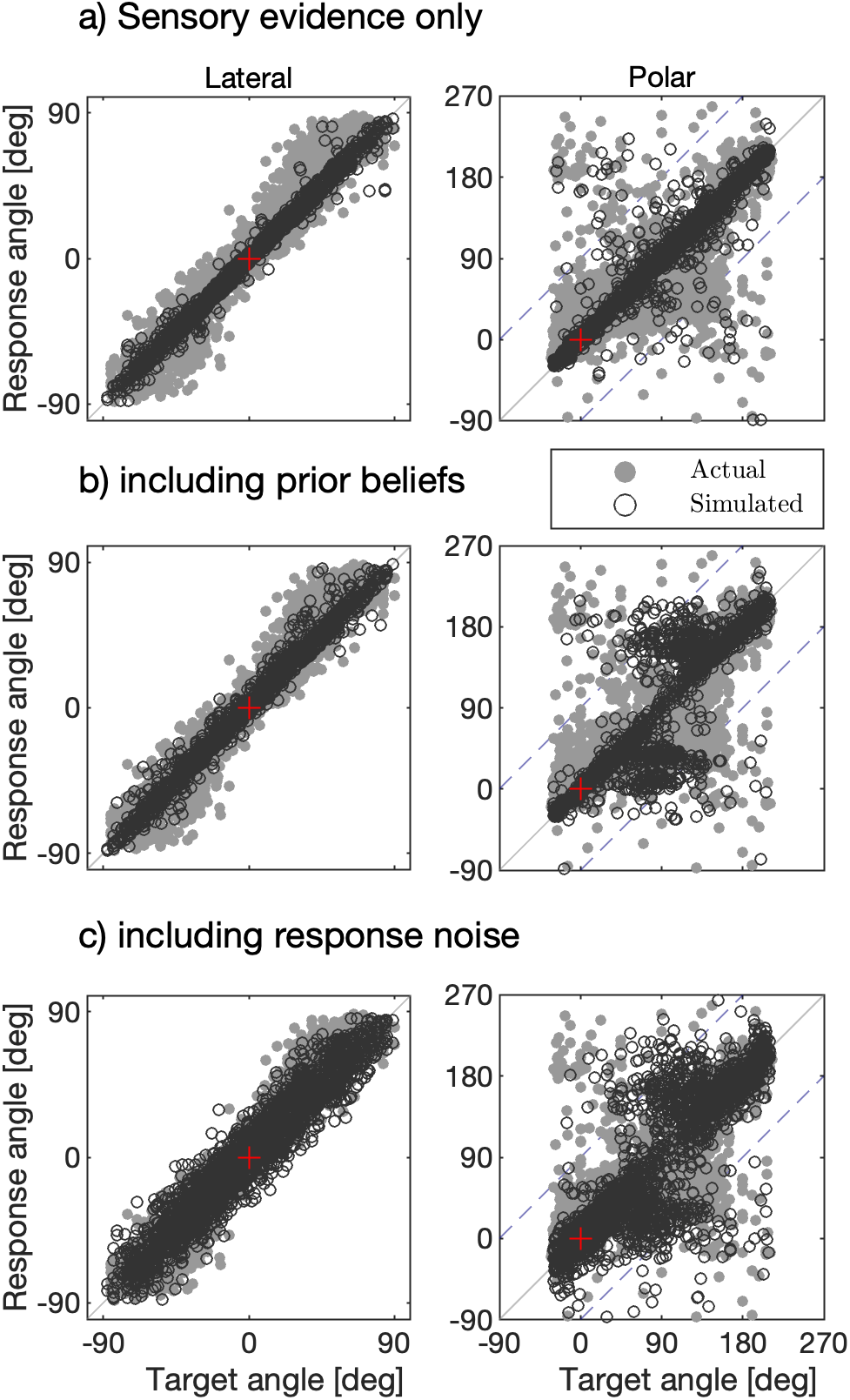
Effects of likelihood, prior, and response noise on predicted response patterns as a result of modeling the directional localization of broadband noise bursts. **a)** Likelihood obtained by sensory evidence (i.e, no spatial prior and no response noise). **b)** Bayesian inference (with the spatial prior but no response noise). **c)** Full model (with prior and response noise). Gray: actual data of NH16 from [28]. Black: estimation obtained by the model considering spectral gradient profiles (GPs). Red cross: frontal position. Blue dashed lines separate regions of front-back confusions.

In order to quantify the effect of introducing the spatial prior in the polar dimension, we computed the polar gain as a measure of accuracy [13] for both simulated and the actual responses. This metric relies on two regressions performed on the baseline condition, separating between targets in the front and back. The linear fits for the baseline condition are defined as:

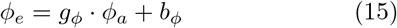

with *ϕ*_*e*_ being the estimated polar angles and *ϕ*_*a*_ being the actual polar angles. The parameters are the localization bias *b*_*ϕ*_ in degrees, which is typically very small, and the dimensionless localization gain *g*_*ϕ*_, which can be seen as a measure of accuracy [8, 13]. The regression fits only incorporate *ϕ*_*e*_ that deviate from the regression line by less than 40°. Since that definition of outliers depends on the regression parameters, this procedure is initialized with *b*_*ϕ*_ = 0°and *g*_*ϕ*_ = 1 and re-iterated until convergence. In our analysis, only the frontal positions were considered. The polar gain of the actual responses, averaged over subjects, was 0.50, indicating that our subjects showed a localization error increasing with the angular distance to the horizontal plane. For the models without the prior, the predicted polar gain was 1.00 (Fig. 6a). The polar gain obtained by the model including the prior was 0.62 (Fig. 6b and c) showing a better correspondence to the actual polar gain. Hence, the introduction of the prior belief improved the agreement with the actual localization responses by biasing them towards the horizon.

### B. Model evaluation

The performance evaluation was done at the group-level. For our model, we used the five calibrated parameter sets with templates ***T*** (***φ***) based on the individuals’ HRTFs as “digital observers”. Group-level results of these digital observers were then evaluated for two psychoacoustic experiments with acoustic stimuli as input that differed from the baseline condition with a flat source spectrum and individual HRTFs.

In addition, we compared our results with the ones of two previously published models. The first one, described by Reijniers et al. [6], is probabilistic and able to jointly estimate the lateral and polar dimensions similar to the model described in this work. Reijniers’ model deviates from the current model since it relies on a different feature extraction stage, uses a uniform spatial prior distribution, does not include response noise (Eq. 13) and does not fit individualized parameters. The second model, described by Baumgartner et al. [15], estimates sound positions only in the polar dimension. Nevertheless, it shares a similar processing pipeline with the current model in that it considers both a perceptually relevant feature extraction stage, includes response noise, and considers individualized parameters. The main differences with our model resides on the incorporation of a directional prior and of the lateral dimension per se, and on how the distance between target and templates is computed. Particularly, this previous work resorted on the *l*^2^-norm to implement the template comparison procedure which is substantially different from our likelihood function. At the moment, this model is commonly used by the scientific community that is interested in elevation perception based on monaural spectral features for sound direction estimation [e.g., 45, 46]. We will refer to these two models as `reijniers2014` and `baumgartner2014`, respectively.

#### 1. Effects of non-individual HRTFs

In first evaluation, sounds were spatialized using non-individualized HRTFs [29]. Originally, eleven listeners localized Gaussian white noise bursts with a duration of 250 ms and sound directions were randomly sampled from the full sphere. Subjects were asked to estimate the direction of sounds that were spatialized using their own HRTFs in addition to sounds that were spatialized using up to 4 HRTFs from other subjects (21 cases in total). With the aim to reproduce these results, we had our pool of five digital listeners localize sounds from all available directions that were spatialized with their own individual HRTFs (*Own*) as well as sounds that were spatialized with HRTFs from the other 4 individuals (*Other*). We thus considered all inter-listener HRTF combinations for the non-individual condition.

Fig. 7 summarizes the results obtained for localization experiments with own and other HRTFs. In the *Own* condition, there is a small deviation between the actual results from [29] and our model predictions. This mismatch reflects the fact that the digital observers represent a different pool of subjects (taken from [28]) tested on a slightly different experimental protocol and setup. Differences in performance metrics are small between the two feature spaces, as already reported during parameter fitting. Predictions from the `baumgartner2014` model are only possible for the polar dimension. Instead, the model `reijniers2014` predicted too small errors, as also observed in previous simulations employing this model [24, 47].

**FIG. 7:**
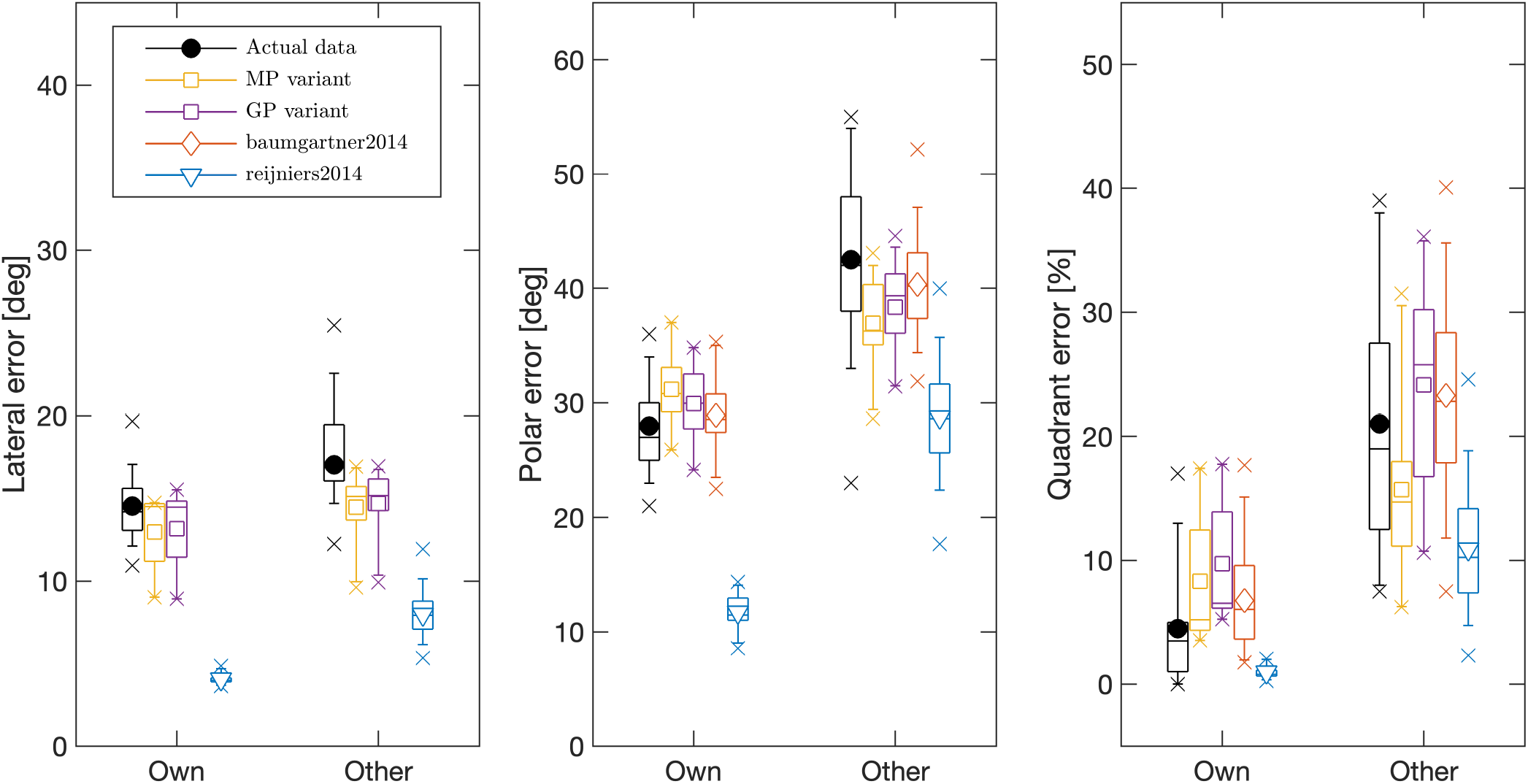
Localization performance with individual (*Own*) and non-individual (*Other*) HRTFs. Actual data from [29] (`data_middlebrooks1999`). Model predictions for two model variants: spectral magnitude profiles (MPs) and spectral gradient profiles (GPs). As references, predictions by the models `reijniers2014` [6] and `baumgartner2014` [15] are shown. Note that `baumgartner2014` does not predict the lateral error.

In the *Other* condition, both of our model variants predicted a smaller degradation for the lateral dimension as compared to the actual data. The lateral errors predicted by `reinjiers2014` increased moderately but remained too small in comparison to the actual data. In the polar dimension, both model variants resulted in increased PEs and QEs, but the amount of increase was larger and more similar to the actual data for the variant equipped with gradients profiles, especially with respect to QE. The predictions from `baumgartner2014` were very similar to the model based on spectral gradients, as expected given the similar method to extract monaural spectral feature. Instead, the simulations form `reijniers2014` demonstrates how this model reported super-human performances as already demonstrated in previous analysis [24].

#### 2. Effects of rippled-spectrum sources

The second evaluation tested the effect of spectral modulation of sound sources on directional localization in the polar dimension [30]. In that study, localization performance was probed by using noises in the frequency band [1, 16] kHz which spectral shape were distorted with a sinusoidal modulation in the log-magnitude domain. The conditions considered different ripple depths, defined as the peak-to-peak difference of the log-spectral magnitude, and ripple densities, defined as the sinusoidal period along the logarithmic frequency scale. The actual experiment tested six trained subjects in a dark, anechoic chamber listening to the stimuli via loudspeakers. The sounds lasted 250 ms and were positioned between lateral angles of ± 30°and polar angles of either 0 ± 60°for the front or 180 ± 60°for the back. A “baseline” condition included a broadband noise without any spectral modulation (ripple depth of 0dB). To quantify the localization performance, we used the polar error rate (PER) as they defined [30]. For every condition, two baseline regressions are computed as in Sec. III A allowing to quantify the PER as the ratio of actual responses deviating by more than 45°from the predicted values of the baseline regression.

Fig. 8 shows the results of testing the fitted models with rippled spectra. In the baseline condition, our model exhibited similar performances to those obtained in the actual experiment, whereas `baumgartner2014` underestimates the baseline performance for this particular error metric. In the ripple conditions, actual listeners demonstrated poorest performance for ripple densities around one ripple per octave and a systematic increase in error rate with increasing ripple depth. These effects were well predicted by the model variant based on gradient profiles, similar to the predictions from `baumgartner2014`. In contrast, both `reijnier2014` and our model based on magnitude profiles were not able to reflect the effects of ripple density and depth as present in the actual data. Hence, the positive gradient extraction appears crucial processing step for predicting sagittal-plane localization of sources with a non-flat spectrum.

**FIG. 8:**
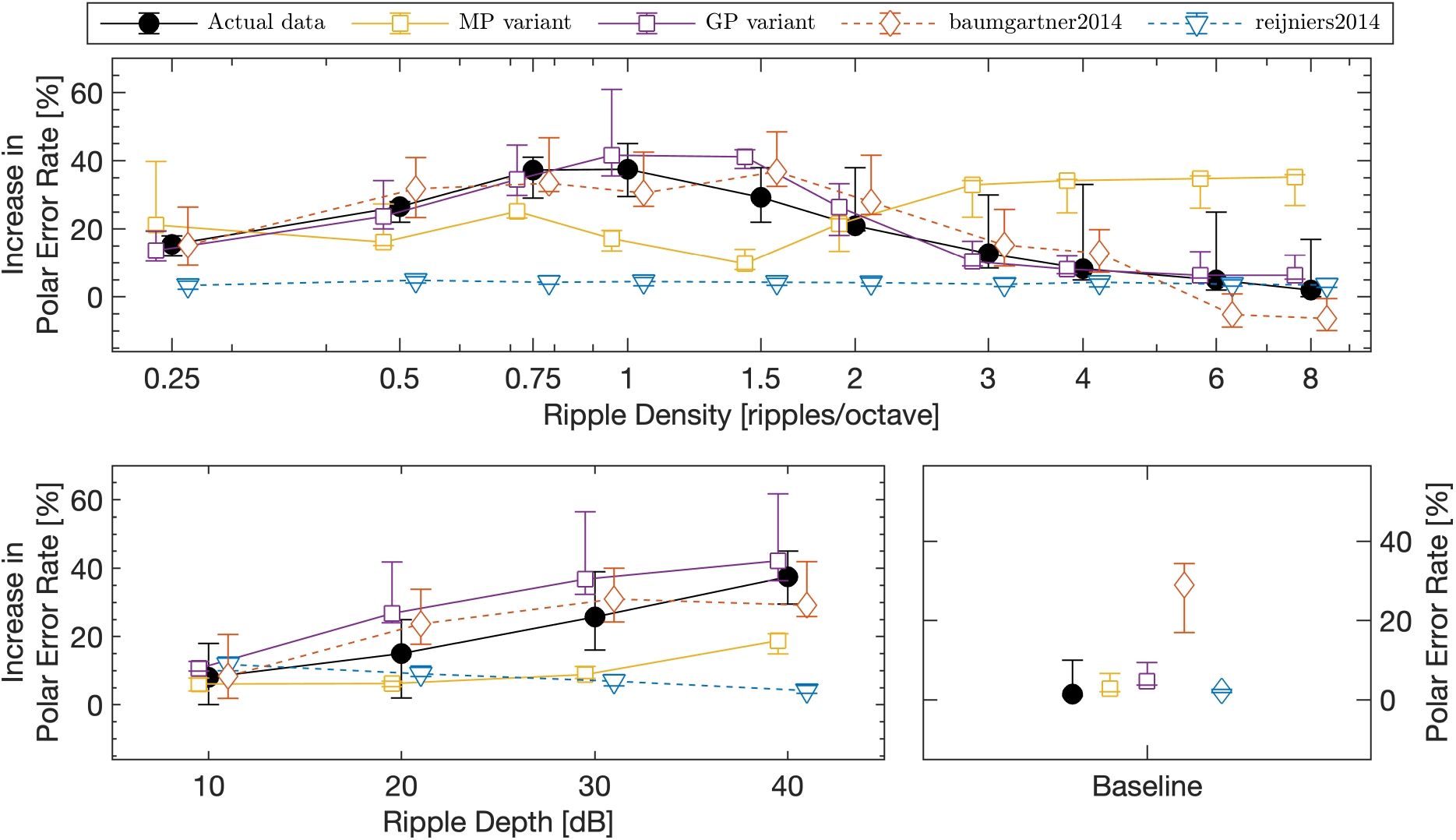
Effect of spectral ripples in the source spectrum on sound localization performance in the median plane. Right-most bottom panel: localization error rates obtained without spectral ripples serving as reference. Top and bottom left panels: Differences to the reference condition shown in the right-most bottom panel. In addition to predictions from the two model variants (MP and GP), predictions from `reijniers2014` [6] and `baumgartner2014` [15] as well as actual data from [30] (`data_macpherson2003`) as shown. Note that ripple densities were varied at a fixed ripple depth of 40 dB and ripple depths were varied at a fixed ripple density of one ripple per octave.

## IV. DISCUSSION

The proposed functional model aims at reproducing listeners’ performances when inferring the sound-source direction.The model formulation relies on Bayesian inference [25] as it integrates the sensory evidence for spatial directions obtained by combining binaural and monaural features [13] with a spatial prior [8]. Our approach considers uncertainties about the various sensory features, as in [6], in addition to the noise introduced by pointing responses [15]. These model components enabled us to successfully match overall performance metrics (LE, PE, and QE) for five subjects (see Tab. II) and within spatially restricted areas (Fig. 5). Importantly, with the inclusion of a spatial prior the model was able to adequately explain listeners’ response biases towards the horizontal plane. Compared to previous models [6, 15], our model better predicted the group-level effects of non-individualized HRTFs and rippled source spectra, yet only if positive spectral gradient profiles 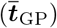, rather than magnitude profiles, served as monaural spectral features.

The validity of the model is limited to scenarios where both the source and subject are spatially static and situated in an acoustic free-field. Additionally, the current model evaluation only considered broadband and stationary sounds. For this reason, we have restricted the extraction of ITDs and ILDs to simple approximations that are sufficient for this set of conditions, as demonstrated both in our present work as well as in previous literature [18, 32] by the high accuracy of estimated lateral angles, where those features are arguable most important. The model, as presented here, does not currently account for feature-specific non-linearities required to predict phenomena like the precedence effect [48, 49] and cannot readily be applied to non-stationary sounds, such as speech, without extensions to the feature extraction procedure. Nevertheless, from an application point of view, the proposed model can be a useful tool to assess the perceptual validity of a non-individual HRTF dataset. As an example, one can use the model’s predictions to quantify differences in expected direction estimation performance for (input) sounds that are spatialized with generic vs. individual HRTFs [50].

In this work, the most probable source position is selected from the posterior distribution via the MAP decision rule. We preferred this widely-used estimator over the equally-common mean estimator to adequately deal with multiple modes of the posterior distribution generated by poor discrimination between front-back positions along sagittal planes. On the other hand, one could argue that the MAP estimator disproportionately biases direction estimates towards the mode where prior probability is large, at least under conditions of high sensory uncertainty. One of many possible alternative decision functions that may better describe stochastic human localization responses is posterior sampling [8]. With this decision function the model would probabilistically sample its best perceptual estimate from the full posterior distribution. Although often considered suboptimal, this strategy would allow an observer faster exploration and flexible adaptation to novel environmental statistics [51]. Importantly, a different decision rule might affect the here-estimated magnitudes of the sensory and motor noise. Therefore, comparative evaluations of different estimators would benefit from a more robust fitting procedure, which falls outside of the scope of the current study.

The model incorporates several non-acoustic components because they are deemed crucial to explain human performances [2, 52]. Extending the `reijniers2014` model [6] by incorporating a spatial prior and response scatter appears vital to explain listeners’ estimation patterns. Without these components, fitting the model to the polar performance metrics was unfeasible [24]. First, response noise allowed us to control response precision at a local level (LE and PE) while leaving global errors (QE) largely unaffected. Instead, the occurrence rate of such global errors depends predominantly on the noise variance that is added to the monaural features. Second, the spatial prior shapes the response patterns by introducing a bias towards the horizon [41]. As shown in Fig. 6, the prior contribution is visible in the polar component of the simulated responses which cluster around the eyelevel direction. Additional evidence is given by the polar gain measure as reported in Sec. III A where the integration of prior beliefs leads to better matching performances in the vertical dimension. Our formulation of the spatial prior was extrapolated from previous work [8] in the sense that its spatial distribution was assumed to be front-back symmetric. Discrepancies observed between actual and predicted global errors (Fig. 5) indicate that this assumption is likely incorrect and points towards an asymmetric prior instead. Nonetheless, at this point we can only speculate about the reasons behind the existence of such a long-term prior in spatial hearing. It potentially reflects the spatial distribution of sound sources during everyday exposure [53], or it may stem from an evolutionary emphasis on high relevance auditory signals [4], or could be related to the center of gaze as observed in barn owls [54] although the processing underlying the spatial inference mechanism might be different in mammals [3]. While the model currently only considers the static scenario, it sets the foundations for future work on predicting sound localization behavior in realistic environments. Evaluating the environment’s dynamics as a chain of consecutive events as in [7] may be a promising approach. Sequential updating of one’s beliefs, from prior to posterior to prior again, comes natural under the Bayesian inference scheme [25]. This makes the here-proposed model well suited as a basis for such investigations. Thus, a rich set of modulators might influence spatial hearing and the model’s prior belief is an entry point to account for many of those, c.f.: evidence accumulation to track source statistics [26, 55], visual influences on auditory spatial perception [56], and auditory attention to segregate sources [57]. Selective temporal integration appears important to deal with the spatial information of many natural sources and their reflections competing in realistic scenarios. This aspect could be partially addressed by integrating recent findings related to interaural feature extraction [58]. To this end, the model will need to consider the dynamic interaction between the listener and the acoustic field. Consequently, these extensions will potentially enable the model to account for subject movements [9] and simultaneous tracking of source movements [59] while extracting spatial information from echoic scenarios [60].

## V. CONCLUSIONS

We proposed a computational auditory model for the perceptual inference of sound-source direction based on Bayesian inference. From a binaural input, interaural and monaural spatial features jointly provide the sensory evidence to estimate the sound direction. The model parameters are interpretable and related to sensory noise, prior uncertainty, and response noise. Having fitted the model parameters to match subject-specific performance in a baseline condition, the model accurately predicted the localization performance observed for test conditions with non-individualized HRTFs and spectrally-modulated source spectra. Regarding spectral monaural feature extraction, the model variant evaluating gradient profiles performed best.

The proposed model seems useful to assess the perceptual validity of HRTFs. The model’s domain is currently limited to static conditions, but it seems to provide a good basis for future extensions to spatially dynamic situations, spectro-temporally dynamic signals like speech and music, and reverberant environments.

## ACKNOWLEDGEMENTS

We thank Jonas Reijniers for providing the initial implementation of his model. We are also grateful to Günther Koliander and Clara Hollomey for helping with the mathematical formulation of some model aspects. This work was supported by the European Union (EU) within the project SONICOM (grant number: 101017743, RIA action of Horizon 2020, to P.M.) and the Austrian Science Fund (FWF) within the project Dynamates (grant number: ZK66, to R.Bau.).

## DATA AVAILABILITY STATEMENT

The implementation of the model presented in this manuscript is available in the Auditory Modeling Toolbox (AMT 1.1) as `barumerli2022` [61]. Data from [28] are also available in the AMT 1.1 as `data_majdak2010`. Individual HRTF datasets of these subjects are publicly available within the ARI database at http://sofacoustics.org/data/database/ari/. Moreover, the implementations of the model `baumgartner2014` presented in [15], the model `reijniers2014` from [6] and the data for the model evaluation procedure are available as `data_middlebrooks1999` and `data_macpherson2003` in the AMT 1.1.

Additional toolboxes were selected for the model implementation. To provide quasi-uniform sphere sampling the model relied on https://github.com/AntonSemechko/S2-Sampling-Toolbox. For the implementation of the von Mises-Fisher distribution, we used https://github.com/TerdikGyorgy/3D-Simulation-Visualization/.

